# 3D Printed Microheater Sensor-Integrated, Drug-Encapsulated Microneedle Patch System for Pain Management

**DOI:** 10.1101/788877

**Authors:** Mengtian Yin, Li Xiao, Qingchang Liu, Sung-Yun Kwon, Yi Zhang, Poonam R Sharma, Li Jin, Xudong Li, Baoxing Xu

**Affiliations:** Department of Mechanical and Aerospace Engineering, University of Virginia, PO Box 400746 122 Engineer’s Way, Charlottesville, VA 22904, USA; Department of Orthopedic Surgery, University of Virginia, 135 Hospital Drive, Charlottesville, VA 22908, USA; Department of Biomedical Engineering, University of Virginia, 135 Hospital Drive, Charlottesville, VA 22908, USA; Theraject, Inc., 39270 Paseo Padre #112, Fremont, CA 94538, USA; Biomedical Imaging Research Institute, Cedars-Sinai Medical Center, 116 N. Robertson Blvd, Pacific Theatres Building, Suite 400, Los Angeles, CA 90048 USA

## Abstract

Microneedle patch device has been widely utilized for transdermal drug delivery in pain management, but is challenged by accurate control of drug release and subsequent diffusion to human body. The recent emerging wearable electronics that could be integrated with microneedle devices offers a facile approach to address such a challenge. Here a 3D printed microheater integrated drug-encapsulated microneedle patch system for drug delivery is presented. The ink solution comprised of polydimethylsiloxane (PDMS) and multiwalled carbon nanotubes (MWCNTs) with mass concentration of up to 45% is prepared and used to print crack-free stretchable microheaters on substrates with a broad range of materials and geometric curves. The adhesion strength of printed microheater on microneedle patch in elevated temperatures are measured to evaluate their integration performance. Assessments of encapsulated drug release into rat’s skin are confirmed by examining degradation of microneedles, skin morphologies, and released fluorescent signals. Results and demonstrations established here creates a new opportunity for developing sensor controlled smart microneedle patch systems by integrating with wearable electronics, potentially useful in clinic and biomedical research.

---

Treatment of low back pain through the systemic administration of medication is an essential approach in pain management, in particular for the pain that is usually associated with a variety of pathological conditions. Transdermal delivery of medication to local on-sites is considered superior to oral administration because of reduced side effects such as peptic ulcer and gastrointestinal bleeding and avoided liver first-pass metabolism.^[1]^ Traditional technologies of drug delivery range from the early topical pain patches to recently developed microneedle patches.^[2]^ Topical pain patches rely on the natural diffusion mechanism of drugs from the surface of the skin into the body, which is limited to the use of small molecular drugs with molecular weight less than 500 Da.^[3]^ In contrast, microneedles that are usually in hundreds of micron length range creates microchannels by penetrating through the skin’s natural defense barrier - stratum corneum (SC) with relatively little or no pain and allows a direct permeation of drugs into the body with enhanced delivery efficiency for a wide scope of drug size and categories, such as peptide, protein, antibodies and nanomaterials.^[4]^ To ensure the skin permeability and achieve the accurate release of drugs that is of critical importance in defining the personalized treatment protocol, microneedles with a range of shapes and materials have been designed, for example, solid microneedles for pre-treatment skin,^[5]^ coated microneedles with medication,^[6]^ hollow microneedles filled with the drug solution,^[7]^ and dissolvable microneedles made of controllable degradable rate of polymer materials.^[8]^ Several physical methods have been developed in conjunction with microneedles to control the diffusion of drugs in the body for disease specific applications of relevance to pain, for example, ultrasound,^[9]^ iontophoresis,^[10]^ electroporation,^[11]^ chemical enhancers.^[12]^

Alternatively, heat that can be taken easily and inexpensively from the form of hot clothes, hot baths, and heating pads/wraps could be used to accelerate the diffusion of drug molecules as the increase of temperature could promote the movement of molecules,^[13]^ thus helping control delivery of drugs. Besides, as a traditional straightforward way, heating has proved to help short-term pain relief such as acute low back pain due to increased blood flow, softened surrounding tissues, and associated metabolism and physiological effects, often referred as heat therapy or thermotherapy.^[14]^ In practical applications, it emerges as an attractive approach in transdermal delivery of medication to integrate the heat therapy with microneedle patches to create a single simple smart sensor-assisted microneedle patch system that could control and monitor the release of drugs in an efficient manner. Recent development of technologies in 3D printing of functional structures and electronic devices could provide an opportunity to manufacture this smart system.^[15]^

Here, we present a drug-encapsulated microneedle patch system integrated with a 3D-printed microheater device and demonstrate that the control of drug delivery rate can be well controlled by regulating the temperature with the microheater. The printing ink solution consisting of multiwalled carbon nanotubes (MWCNTs) and polydimethylsiloxane (PDMS) is developed with the concentration of MWCNTs up to 45% (10 time higher than ever)^[16]^ and the 3D printing ability of microheater on various material and geometric types of substrates are demonstrated. In particular, its adhesion with the surface of carboxymethyl cellulose enabled microneedles is evaluated as the heating function operates, and a facile strategy is proposed to enhance their integration adhesion by tuning the ink solution. *In vitro* tests on rat’s skin is conducted to assess the functional abilities of the microheater integrated microneedle patch system in drug delivery. These studies provide a new route to design and manufacturing of sensor controlled medical devices and also establish the foundations for improving pain management via microneedle patches by seamlessly integrating wearable electronics to meet requirements for clinical use.

## Materials, design and 3D printing of microheater

We first investigate the droplet spreading of the MWCNTs/PDMS ink solution on a solid surface. **Figure 1a** shows the schematic illustration of an ink droplet with the contact diameter of *R*_*d*_ printed on a curved substrate with the curvature of *R*_*t*_. Heights of the droplet before and after spreading are denoted as *H*_*i*_ and *H*_*s*_, respectively. The system total energy of the droplet on the surface at equlibrium that includes both the solid-liquid interaction energy and gravitational potential energy of the droplet proves to be a functional of *H*_*s*_*/H*_*i*_. With the energy minimization and the conservation of droplet mass, *H*_*s*_*/H*_*i*_ can be quantitatively determined and will depend on the radius of printing needle and substrate, *R*_*nozzle*_ and *R*_*t*_ (See method for theoretical analysis, and **Figure S1 in Supporting Information**). Here we define the liquid spreading will not occur when *H*_*s*_*/H*_*i*_ is larger than 90%, i.e. the final collapsed height is more than 90% of the initial height, and the printed devices on this curved substrate will achieve the pattern as designed. **Figure 1b** shows a theoretical map of a liquid drop on a curved substrate. With the definition of droplet spreading via *H*_*s*_*/H*_*i*_, spreading and non-spreading regions with respect to the nozzle size and curvature of substrate can be obtained, and it will change with surface tension of droplet ink (i.e. concentration of MWCNTs in ink solution). In general, larger curvatures of substrates and ink droplets or lower concentrations of MWCNTs will lead to easier spreading of extruded ink droplets. Multiwalled carbon nanotubes (MWCNTs) and polydimethylsiloxane (PDMS) with 10:1 of base polymer to curing agent were used to prepare the solution inks ^[17]^, and the extrusion-based ink jet printing (Sterile standard blunt needle, 1.8mm, Cellink, Inc.) was used to generate an ink droplet. **Figure 1c** shows the optical images of the MWCNTs/PDMS ink droplet with MWCNTs concentration of 10% on a flat PDMS substrate as printed and after 30 minutes. The measured *H*_*s*_*/H*_*i*_ is 0.92, indicating there is no spreading of the droplet after equilibrium. By contrast, when the concentration of MWCNTs decreases to 3%, *H*_*s*_*/H*_*i*_ will reduce to 0.423, and a clear spreading is observed (**Figure S2a in Supporting Information**). **Figure 1d** and **Figure S2b in Supporting Information** shows optical images of the ink drop printed on curved surface with the curvature radius of 20 mm as printed and after 30 minutes. *H*_*s*_*/H*_*i*_ is 0.938 and 0.594 for the MWCNTs concentration of 10% and 3%, respectively, which corresponds to non-spreading and spreading of the droplet on the surface, similar to those observation in the flat substrate. Both measurements with these two different MWCNTs concentration in the ink solutions show good agreement with theoretical predictions. Besides, a higher *H*_*s*_*/H*_*i*_ is obtained on a curved substrate, which is in good consistence with theory as well.

**Fig 1.**
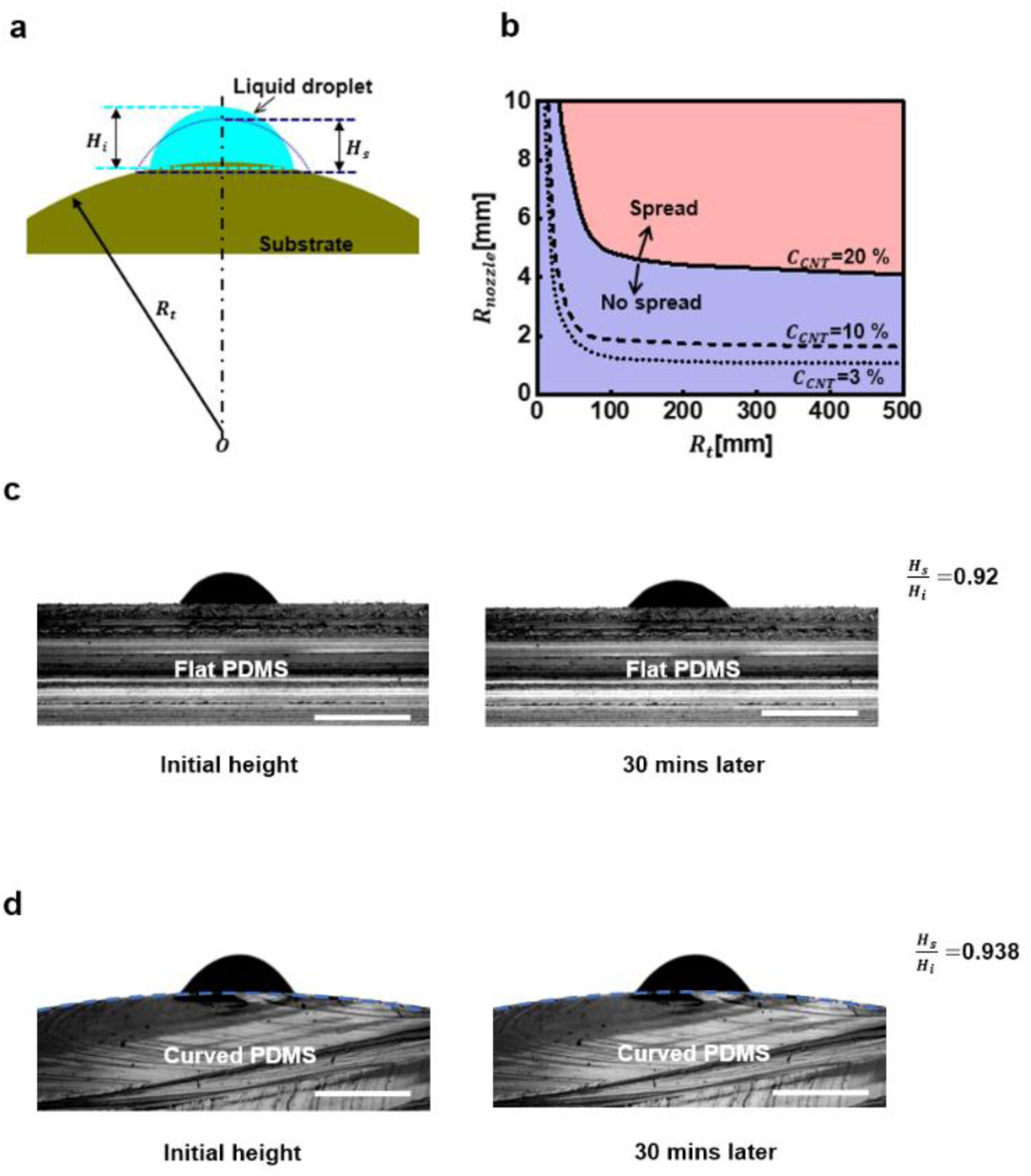
Theoretical model and experiment of the 3D printed droplet spreading on a curved substrate. **(a)** Schematic illustration of a 3D printed liquid droplet on a curved substrate. The radius of curvature of the substrate is *R*_*t*_, and heights of the droplet before and after spreading are *H*_*i*_ and *H*_*s*_, respectively. **(b)** Theoretical map of a 3D printed liquid droplet spreading on a curved substrate in the variation of *R*_*t*_ and nozzle size of 3D printer *R*_*nozzle*_. The droplet is defined to spread at *H*_*s*_/*H*_*i*_<90% and otherwise will not. **(c)** Optical images of the MWCNTs/PDMS droplet (10% concentration of MWCNTs) on the flat PDMS substrate for as-printed and after 30-minute status. *H*_*s*_/*H*_*i*_ = 0.92 indicates no spreading of droplet. **(d)** Optical images of the MWCNTs/PDMS droplet (10% concentration of MWCNTs) on the curved PDMS substrate (curve radius: 20 mm) for as-printed and after 30-minute status. *H*_*s*_/*H*_*i*_ = 0.92 indicates no spreading of droplet. *H*_*s*_/*H*_*i*_ = 0.938 indicates no spreading of droplet. All droplets in (c) and (d) were printed with the diameter of 18-gauge print head and were in 1.8 mm in diameter as printed. Scare bar for (c) and (d) is 2 mm.

With the fundamental elucidation of ink droplets on the substrates, we will print the microheater with the same MWCNTs/PDMS ink solution (see method). Hexane was added to decrease the viscosity of ink solution and benefit the extrusion from nozzles, which is of critical importance for a high concentration of MWCNTs (>5%).^[18]^ The extrusion rate of ink solution was 12 mm/s and the thickness of printed layer 800 μm. The printed devices were cured in 80°C for 2.0 hours. **Figure 2** shows optical photographs of the as-printed microheater in a meander pattern with dimensions of 30 mm (length) × 20 mm (width) (**Supplementary movies**). The meander pattern consists of six turns with a gap of 2.5 mm, measured from center to center. Two squares at upper left corner and lower right corner are 5 mm × 5 mm. Six different printing substrates were chosen, including a flat surface of paper, PDMS and glass, and microneedle patch and a curved surface of apple skin and gloves represent, and no spreading of the ink or destruction of the pattern is observed. Besides, when the paper and PDMS substrates were bended or twisted, neither cracks in the pattern nor delamination between pattern and substrate are observed in microheater (**Figure S3 in Supporting Information**), demonstrating the flexibility of printed pattern and ink solution after cure. The same observations are also seen for the printed microheater in a double spiral pattern (**Figure S4 in Supporting Information**), further indicating the robustness of printing process and preparation of ink solution. The double spiral pattern is 30 mm in length and width, with three resolutions. The gap between each line is 3 mm, from center to center. We should note that when the concentration of MWCNTs in ink solution was larger than 25%, onetime cure in a constant temperature of 80°C would lead to cracking in devices due to rapid release of printing residual stress associated with bubbling. A post-annealing treatment is proposed to cure the printed devices to eliminate cracking, as shown in **Figure S5 in Supporting Information**.

**Fig 2.**
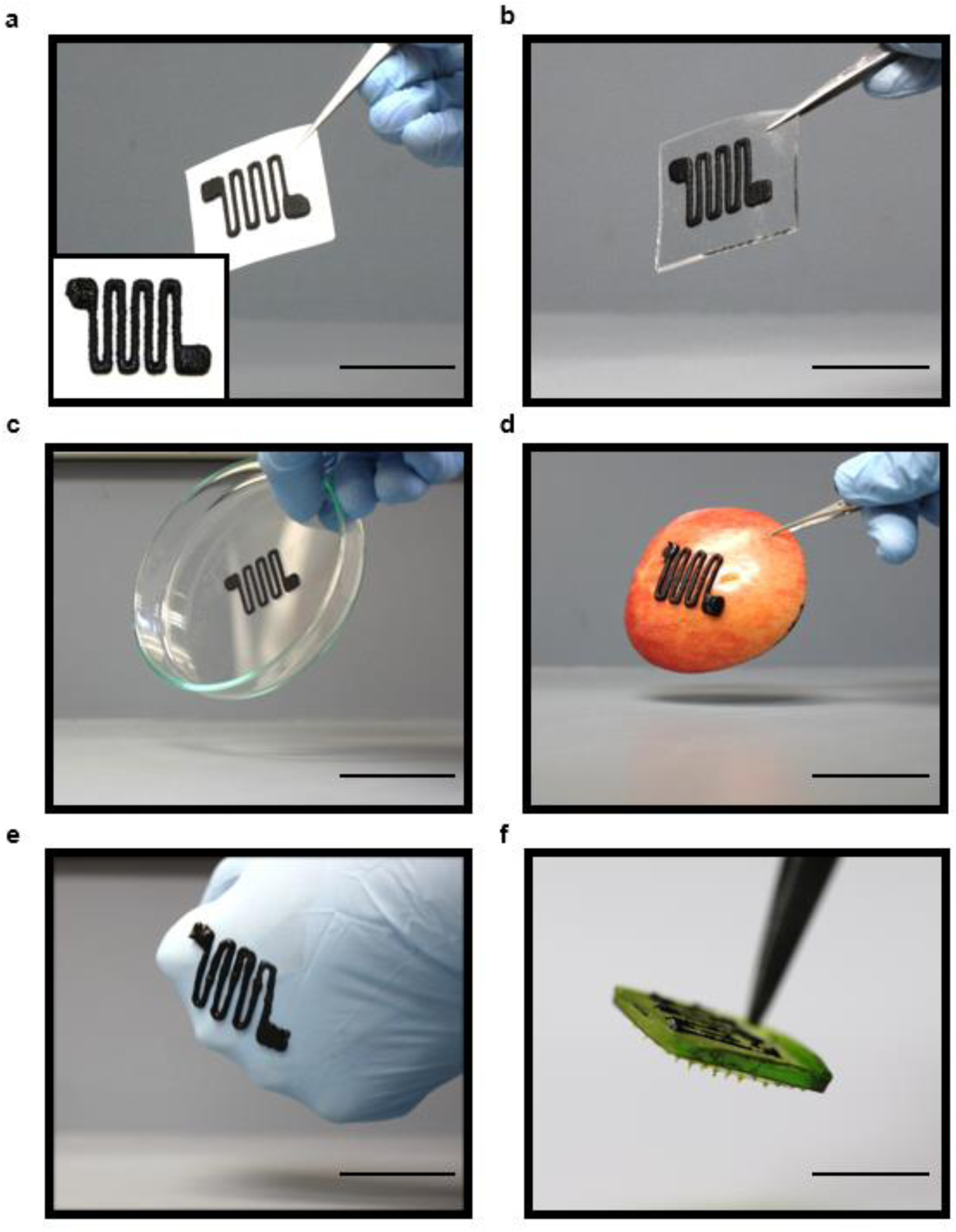
Optical images of microheaters printed on different substrates. **(a)** Printer paper. **(b)** PDMS. **(c)** Glass Petri dish. **(d)** Curved surface of apple. **(e)** Hand with experimental glove. (f) Top surface of microneedle patch (cellulose). Scale bar is 35 mm for (a)-(e), and 4.5 mm for (f).

## Fundamental studies of microheater and integration with microneedle patch

To test the heating performance of the printed microheater, a temperature sensor was attached on its top surface and placed at room temperature (∼20°C). A power supply is connected to both ends of the microheater, and with the current flowing in conductive composite, heat is generated and conducted to patch, microneedle, and skins (**Figure S6 in Supporting Information**). Take the printed microheater with 20% concentration of MWCNTs on PDMS substrate as an example, upon applying a constant voltage, **Figure 3**a shows that the temperature increases at the beginning and reaches equilibrium state due to an open environment to air. With the increase of applied voltage, a higher equilibrium temperature is obtained with almost linear proportional to the rise of applied power, suggesting the typical joule heating mechanism in the microheater. **Figure 3b** shows the measured temperature change under a cyclic loading condition of voltage. The same of equilibrium temperature without obvious thermal degradation in each voltage cycle indicates the stability of heating performance of the printed devices for a long-term use. Figure 3c further presents the effect of the concentration of MWCNTs in ink solution on the electrical resistance and equilibrium temperature in printed microheaters. The electrical resistance shows a dramatic drop when the concentration of MWCNTs increases from 20% to 35% and decreases in a relatively slow manner beyond 35%, contributed by the formation of intense conductive networking in PDMS. As a consequence, the equilibrium temperature shows an increase at a higher concentration of MWCNTs. Figure 3d shows the variation of electrical conductance in the devices on PDMS substrate under a stretching loading. PDMS substrate is stretched with a homemade stretcher, as inset at the upper left corner. Approximately linear relationship between them is obtained, independent of the concentration of MWCNTs. The sudden increase of electrical conductance at ∼30% stretching strain suggests the failure of microheater. This large stretchability could easily coordinate with large deformation (∼20%)^[19]^ of human skin accompanied with typical physical activities, potentially useful for pain relief by printing microheater on the skin.

**Fig 3.**
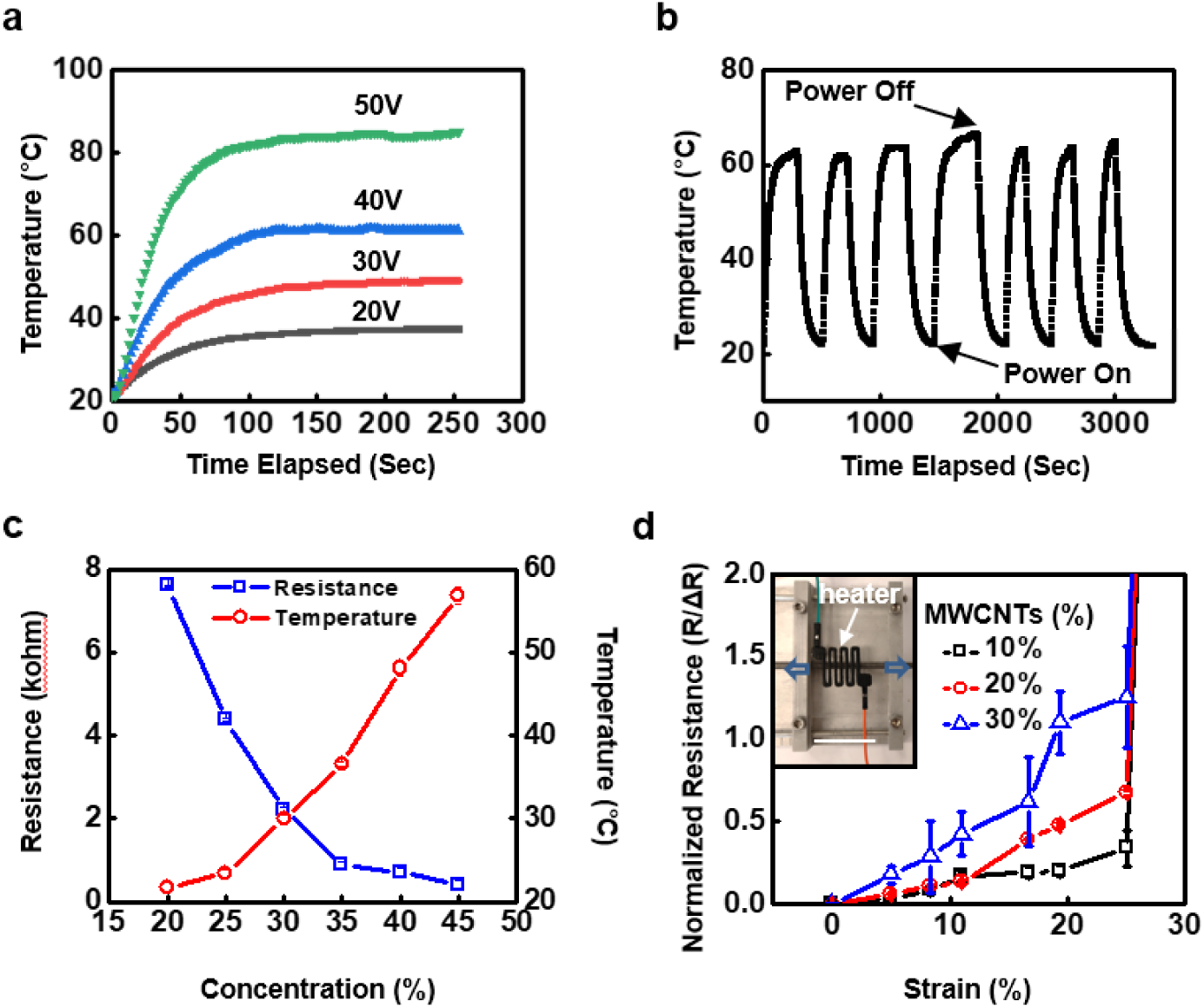
Electrical and mechanical properties of microheater printed on PDMS substrate. **(a)** Measured temperatures of microheater under power supply of 20V 30V 40V and 50V. **(b)** Repeatability test of heating function by microheater. **(c)** Effect of MWCNTs concentration in ink solution on electrical and heating functions of printed microheaters. **(d)** Normalized electrical resistance of microheater subjected to a stretching strain. Inset shows the mechanical stretcher with microheater loaded. The concentration of MWCNTs in ink solution was 20% for and (b). Scale bar for inset is 30 mm.

When microheater is printed on the back surface of microneedle patch (Figure 2f), we performed peel-off experiments to investigate their adhesion strength under an elevated temperature. **Figure 4**a shows the peeling force-displacement curves under a various equilibrium temperature generated by microheater. Typical three stages are observed, 0–2 mm and 4–6 mm for delamination between PDMS layer and cellulose microneedle patch substrate, and 2-4 mm for detachment of microheater from the cellulose microneedle patch substrate. In addition, a higher heating temperature leads to a lower peeling force. It is noteworthy that the slight increase of peeling force at peeling distance of 2-4 mm results from the meander pattern of microheater. Further, we performed the peel-off experiments on printed devices with different ratios of base polymer to curing agent of PDMS in the preparation of PDMS-MWCNTs ink solution. **Figure 4b** shows the effect of PDMS in ink solution on adhesion strength at the heating temperature of 44°C. A lower concentration of PDMS curing agent in ink solution leads to a higher peeling force, suggesting a subtle approach to prevent detachment of microheater from microneedle patch at elevated heating temperatures.

**Fig 4.**
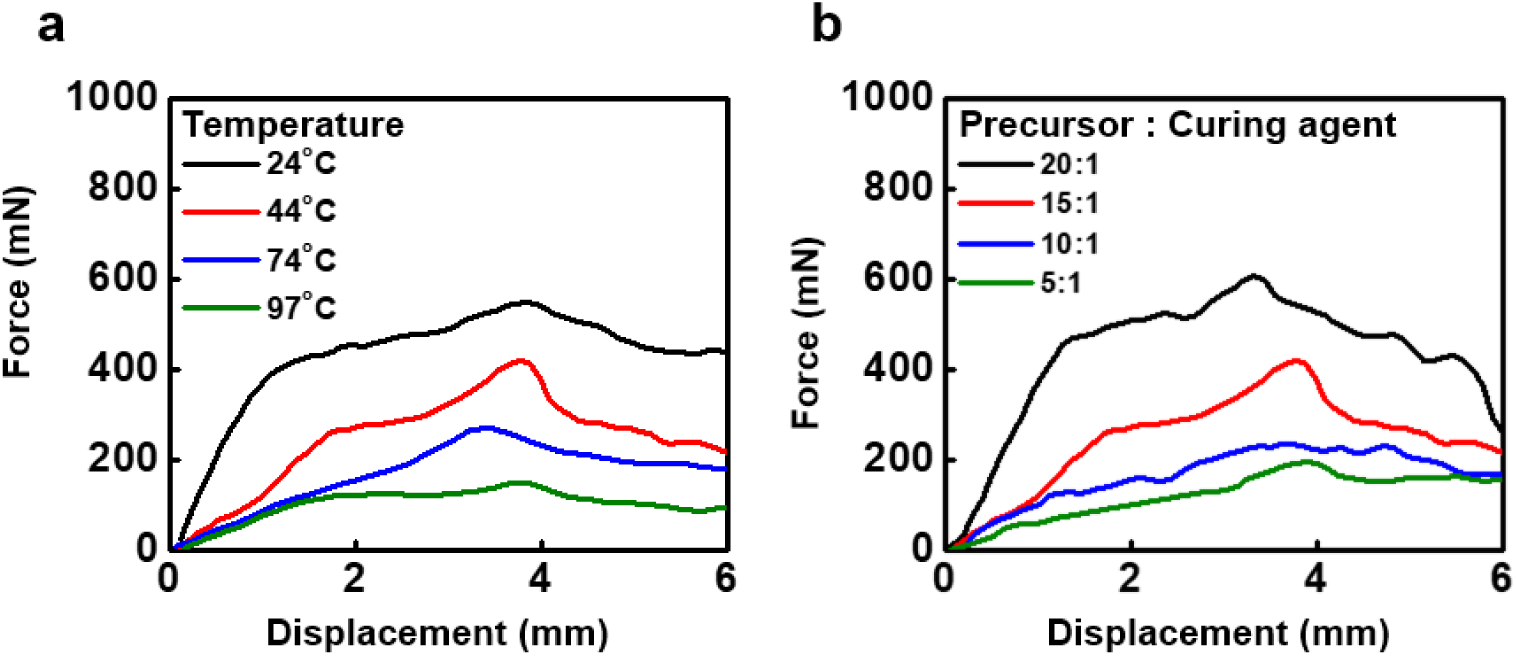
Adhesion performance of PDMS/printed microheater on microneedle system (8% cellulose). **(a)** Peel-off force – displacement of the system with PDMS base/curing agent of 15:1 in prepared ink solution at the heating temperature of 24°C, 44°C, 74°C, 97°C. **(b)** Peel-off force – displacement of the system with different ratios of PDMS base to curing agent in the prepared ink solution under heating temperature of 44°C.

## *In vitro* evaluations using rat skin

Evaluation of drug diffusion and release using freshly excised skin specimen provides a viable approach for validating physiochemical performance of microheater integrated microneedles. The microheater with the MWCNTs concentration of 20% in the ink solution was directly printed on the back surface of microneedle encapsulated with a model drug—the near-infrared fluorescence dye Cy5 (Figure 1f). After the power is switched on, microheater starts to generate heat at desired temperature to facilitate microneedle drug delivery from three aspects: 1) Microheater accelerates the dissolving process of microneedles in the presence of skin fluid (when microneedles inserted into the skin) and thereby accelerating the release of encapsulated drug molecules. 2) Heat or mildly elevated skin temperature can facilitate drug diffusion from the stratum corneum/epidermis (the microneedle application site to deeper dermis tissues that are rich in nerves and blood vessels, due to increased thermal movement of drug molecules. Reaching the dermis tissue in a faster mode ensures the faster-acting therapeutic effect of drug. This “fast-acting” characteristic is particular important when treating pain symptoms. This is also the reason why “heated lidocaine pain patch” has been widely studied and utilized in the clinical setting to alleviate pain. 3) It is well known that local heat can increase blood circulation which further facilitate the drug diffusion and absorption to improve the treatment outcome. We did not observe any skin damage or irritation caused by the heating as showed in **Figure S7 in the Supporting Information**. **Figure 5**a shows the comparison of microneedles before being inserted into skin and after stay inside skin for 10 mins with and without integration of microheater. Microheater clearly facilitated the microneedle dissolving process in rat skin as shown in diminished shape and dimension of microneedle materials, which was accompanied by accelerated release and diffusion of encapsulated drug Cy5 into skin after inserted into skin. **Figure 5b** presents the representative optical images of rat’s skin immediately after a 10-min application with microneedle patch. Structural changes in skin tissue with a microneedle shaped pores underneath the microneedle application site are observed, confirming successful insertion of microneedle patch into the skin, which is further corroborated by the histological image (hematoxylin/eosin staining) of the microneedle insertion site in rat skin (**Figure S8 in Supporting Information**). It is very interesting to detect that the microneedle with microheater group showed visible and darker Cy5 (bluish) marks on the application site, indicating more drug loading in the skin due to an elevated temperature, which resonates accelerated microneedle dissolving in Figure 5a. **Figure 5c** suggests more Cy5 associated fluorescent signal could be detected in microneedle with microheater group in skin compared to the microneedle only group. The deeper and wider distribution and stronger intensity of fluorescence signal in red further indicates the accelerated release and diffusion of the model drug Cy5 into skin due to the heating function of the microheater. Note that the intensity of Cy5 is used as an index instead of total amount delivered, due to the altered inter-molecular interaction induced fluorescence quenching, when Cy5 or any fluorescence dye is present in a higher concentration.

**Fig 5.**
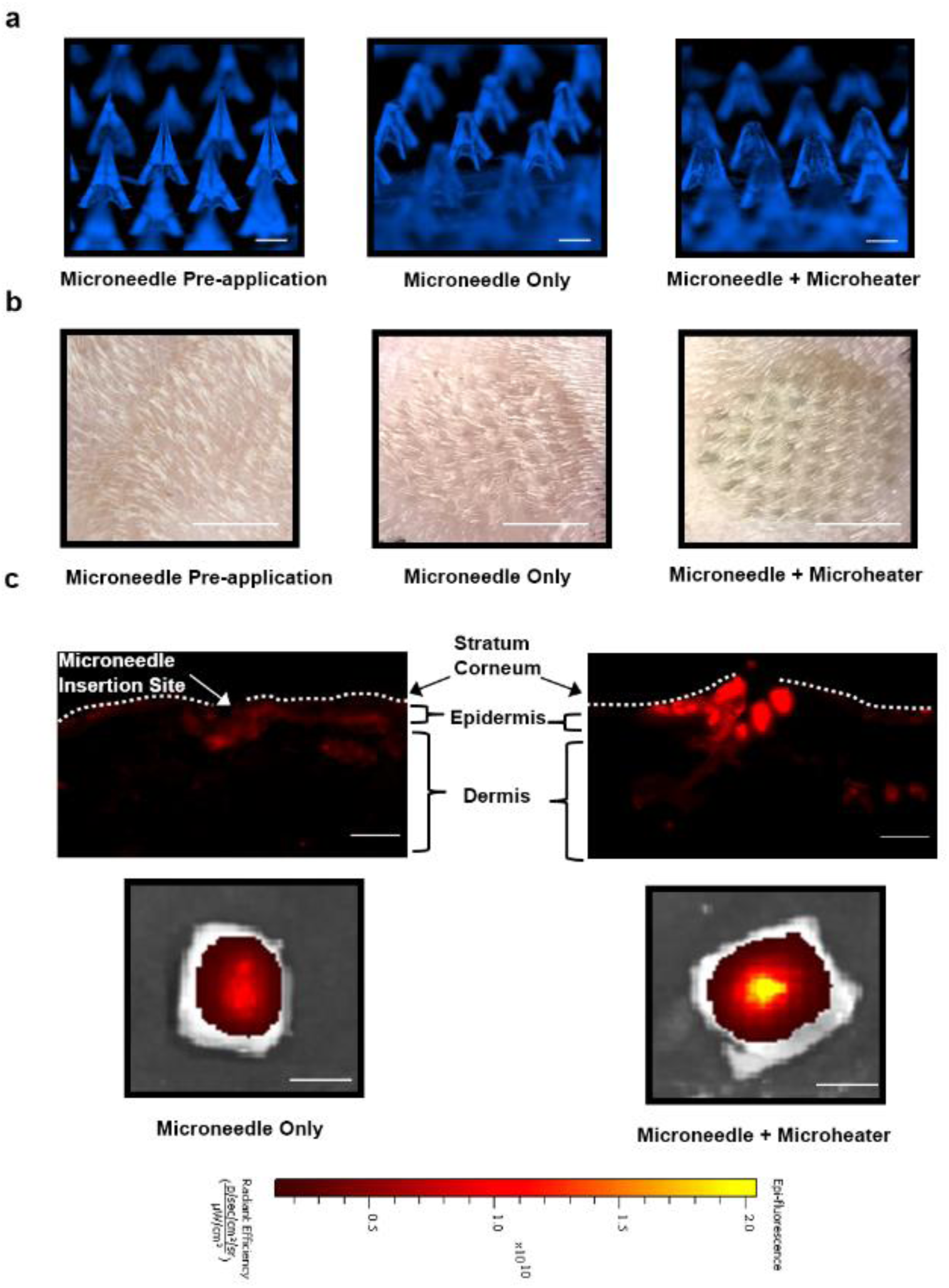
*In vitro* assessment of microheater/microneedle integrated system in control release of a model drug Cy5 in rat skin. **(a)** Optical images of as-fabricated microneedles (pre-application) and microneedles without and with microheater function (at 44°C) after insertion into rat skin after 10 mins. **(b)** Representative pictures exhibited skin marks left by microneedle application confirming successful insertion in rat skin. **(c)** Representative fluorescence images of rat skin tissue slides indicated wider and deeper diffusion of model drug Cy5 (upper row) and more initial drug depot in skin (lower row) in microheater integrated group compared to microneedle only group. Scale bar is 500 μm for (a), 5 mm for (b), 100 μm for first two in (c) and 5 mm for last two for (c).

The therapeutic performance of microneedles and other topical administration methods could be assessed by the amount of initial drug loading on-site as well as the diffusion of drug molecules in the blood circulation through the skin. We adopted an established *in vitro* drug release model to assess the diffusion property of our system using rat skin. **Figure 6**a demonstrates accelerated model drug Cy5 release from rat skin administered with the microheater assisted microneedle compared to microneedle only group. By comparing the quantitative fluorescence intensity of Cy5 diffused from post-application rat skin (with the same 1 ×1 cm^2^ dimension encompassing the entire microneedle application areas), we observed that microheater facilitated more and faster drug release and diffusion (higher fluorescence intensity) along the time for up to 48 hr post incubation of skin specimen in PBS buffer (**Figure 6b**). The slow release phase of the first 25 min followed by a much faster rate in Figure 6 can be interpreted as the traveling time of model Cy5 drug to reach the contact surface of skin and PBS aqueous buffer. This result is consistent with the observation in Figure 4c, indicating that our microheater integrated microneedle system not only increases the initial drug depot at local skin application site but also facilitates drug release and diffusion via accelerating microneedle dissolving, drug release and drug permeation depth in skin.

**Fig 6.**
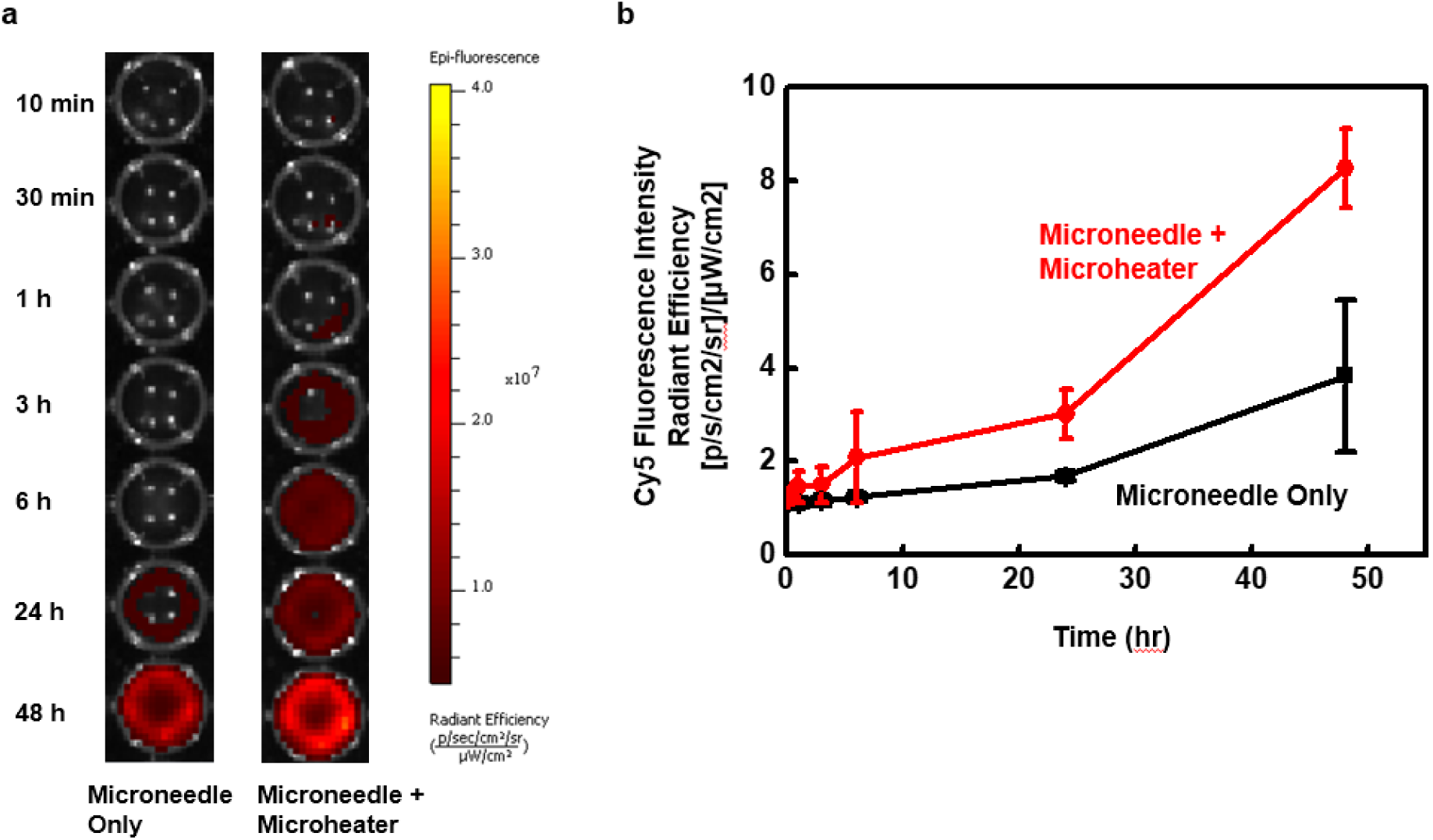
*In vitro* diffusion assay of fluorescence model drug Cy5 release from rat skin after 10-min application microheater/microneedle integrated system. **(a)** NIRF images depicts time-dependent release of Cy5 fluorescent dye in PBS buffer from floating rat skin specimen. **(b)** Quantitative analysis of fluorescence intensities at various time points post incubation in PBS confirmed the microheater promoted drug diffusion.

In summary, we reported a microheater integrated microneedle patch system for drug release and diffusion control that can address the critical needs in pain management with on-demand dose requirement. The preparation of high concentration MWCNTs/PDMS ink solution that can be directly utilized for 3D printing devices including microheater in substrates with a broad range of material and geometric features lays the foundation for the development of ink-based electronics devices with capability of sustaining large deformation and flexibility. The integration performance with tunable interfacial adhesion strength by adjusting PDMS curing ratio in ink solution indicates a facile approach to achieve a robustness interface, even at elevated temperature conditions. *In vitro* evaluation experiments to track the release of drug from microneedle into rat’s skin demonstrate the controlling ability of drug release by heating function via integrated microheater. These results pave a new route to develop electronic sensors integrated controllable smart drug delivery system with broad potential applications of drug molecules in disease treatments.

## Experimental Section

### Theoretical model and analysis of a 3D printed ink droplet on curved substrate

Consider the 3D printed liquid droplet riding on a curved substrate with a radius of curvature *R*_*t*_ (Figure S1a), before the spreading occurs, the height of the droplet is

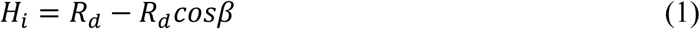

where *β* = *θ*_*c*_ + *α, α* = *asin*(*R*_*c*_/*R*_*t*_), and *R*_*d*_*=R*_*c*_/*sin*(*β*). *θ*_*c*_ is the intrinsic contact angle of the liquid droplet on the corresponding flat surface and *R*_*c*_ is the contact radius. The total surface energy of the system is

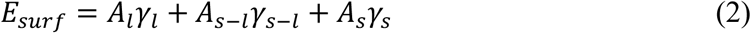

where *A*_*l*_ is the surface area of the liquid, *A*_*s*−*l*_ is the interface area between the substrate and liquid, and *A*_*s*_ is the area of the substrate exposed to the air. *A*_*l*_ = 2*πR*_*d*_*H*_*i*_, *A*_*s*−*l*_ = 2*πR*_*t*_*H*_*t*_, and *H*_*t*_*=R*_*t*_ − *R*_*t*_*cosα* is the height of the spherical cap of the substrate covered by the liquid droplet. *γ*_*l*_ is the surface tension of liquid, *γ*_*s*−*l*_ is the interfacial surface energy density between the substrate and liquid, and *γ*_*s*_ is the surface energy density of the substrate. With Young’s equation, we will have

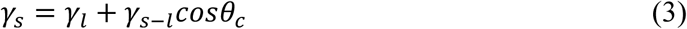

On the other hand, define the center of substrate (*O*) as the reference point, the gravitational potential energy of the droplet can be obtained via

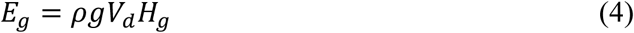

where *ρ* is the density of liquid, *g*(=9.8 m/s^2^) is the acceleration of gravity, 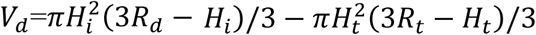is the volume of liquid, and 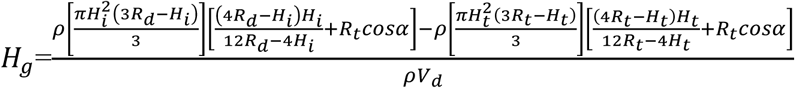 is the position of the center of mass of the droplet. Therefore, the total energy of the system is

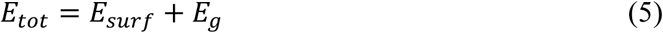

When the spreading of the liquid droplet occurs (Figure S1b), assume the height of droplet changes from *H*_*i*_ to *H*_*s*_(< *H*_*i*_), and the contact angle changes from *θ*_*c*_ to *θ*_*c*_′ (< *θ*_*c*_), the contact radius and radius of curvature of droplet change from *R*_*c*_ and *R*_*d*_ to 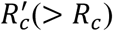 and 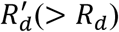, respectively. Similar to Eq. (1), we will have

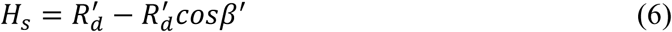

where 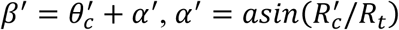and 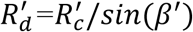.

The surface energy of the system becomes

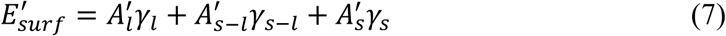

where 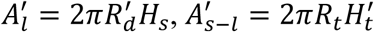, and 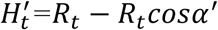. And the gravitational potential energy becomes

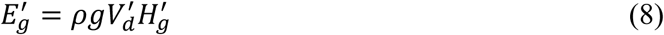

where 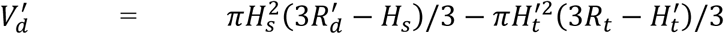, and 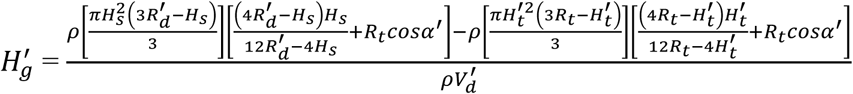. Therefore, the total energy of the system becomes

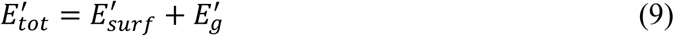

The conservation of liquid volume 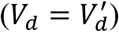 leads to

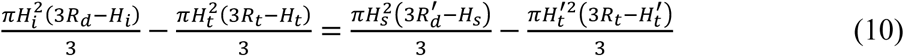

Besides, assume the 3D printed liquid droplet is spherical, we will have 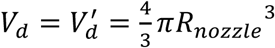, where *R*_*nozzle*_ is the radius of the ink droplet.

Based on Eqs. (9) and (10), we can solve 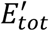 as a function of *H*_*s*_. When 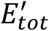 reaches its minimum, the corresponding *H*_*s*_ is the height of droplet after spreading. We here define if *H*_*s*_/*H*_*i*_ ≥0.9, the change of height after spreading is very small and the droplet spreading can be neglected. Otherwise, if *H*_*s*_/*H*_*i*_ <0.9, the droplet spreading will happen. Figure S1c shows the variation of the total energy after spreading normalized by the total energy before spreading 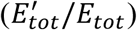 with the height of the droplet after spreading over the height of droplet before spreading (*H*_*s*_*/H*_*i*_). When the total energy reaches the minimum, *H*_*s*_*/H*_*i*_ can be determined as marked. The larger droplet (i.e. a large *R*_*nozzle*_) is, the smaller *H*_*s*_*/H*_*i*_ will be, suggesting the easier spreading of droplet. Similar findings are also found at a lower concentration (i.e. lower surface tension of liquid) of MWCNTs in MWCNTs/PDMS ink solution (Figure S1d).

### Preparing of ink solution and 3D printing of devices

PDMS pre-cursor (Sylgard 184 Silicone Elastomer, Dow Corning Corp.) was firstly mixed with hexane solution (n-Hexane, anhydrous, 95%, Sigma-Aldrich) in a weight ratio of 10:4 in a 50 ml conical tube. Hexane was added to the mixture to ensure a moderate viscosity of mixture. Then, MWCNTs (8-15 nm diameter, 10-15 μm length, 95%, Nanostructured & Amorphous Materials, Inc.) was added to the mixture in a certain concentration (MWCNTs: PDMS in weight). After robust stirring with a stir bar, cartridge was delivered to an ultrasonicator for at least 12 hours with 40 kHz to achieve a homogeneous distribution of MWCNTs in PDMS-Hexane mixture. The mixture was then degassed with a vacuum chamber. Afterward, curing agent of PDMS was added to the mixture in a ratio of 10:1 by weight. The prepared ink solution was loaded to 3D printer (Cellink, Inc.) for use. After several printing, the optimized size of printhead for this research was set to be 14G to decrease the risk of clogging meanwhile keeping the width of microheater within tolerance.

In the printing of devices, the inks were extruded with a printing speed of 12 mm·s^−1^. Printing layer height was 800 μm. After the desired pattern was printed, it was delivered to a controlled temperature chamber to be cured for 2 hr at 80°C. When the concentration of MWCNTs is as high as >25%, to avoid cracking, printed devices were first cured in temperature of 50°C for 10 min and then for room temperature of 25°C for another 10 min, which will help release of printing residual stress and air bubble in devices. This heating-cooling cycle, often referred to annealing process, was repeated for several times till the devices were completely cured.

### Fabrication of microneedle patch

The fabrication of microneedle was conducted by following the protocol published in this paper.^[20]^ In brief, ∼2 mg Cy5 (gift from Dr. Zhang lab at Cedars-Sinai, CA) was dissolved in 500 μL 8% (w/w) sodium carboxylmethyl cellulose (SCMC, Sigma-Aldrich, MO, USA) solution. Then 50 μL of this solution was added to a polydimethylsiloxane (PDMS, Sylgard 184, Dow Corning, USA) mold containing cone cavities (44 cone cavities, each 1 mm in depth and ∼440 μm in base diameter; TheraJect, Inc., CA, USA), and centrifuged at 4000 rpm for 5 min to drive the solution into the cavities. The solution out of the cavities was removed after centrifugation and collected for reuse. After air-drying overnight, another 200 μL solution containing 8% (w/w) SCMC alone was poured on the mold and centrifuged at 4000 rpm for 1 min. The solution was dried overnight to form a microneedle patch. Then patch was peeled off from the mold with caution. After fabrication, microneedles patches were stored at 4 °C until use.

### Biological evaluation using rat skin

Dorsal skin (12-14 wk, male) was obtained from cadaver Lewis rats. Skin hair was removed by hair clipper to provide a clean application area of ∼3 cm×3 cm immediately after rats were euthanized in CO2 chamber. Microneedle with or without heater were then applied on rat skin for 10 min at either room temp (w/o microheater) or at 44°C (by microheater) in designated area on a foam box platform with four corners of skin extended by needle pinching. Optical images of skin before and after microneedle applications were taken to record the gross condition of skin surface and to confirm microneedle insertion. After microneedle application, skin areas subjected to microneedle application were dissected in ∼1 × 1cm^2^. The near-infrared fluorescence (NIRF) images of rat skin specimen after microneedle application were obtained to further confirm the successful microneedle insertion into the skin and to evaluate the amount of model drug deposited immediately after application. Meanwhile, microneedles (both pre- and post-application) were attached in a 45° angel to glass slide with a clear tape and imaged with a Nikon DS-Ri2 digital microscope camera mounted on a Nikon Eclipse E-600 microscope with a DAPI (UV) channel to assess the gross structural change of microneedles with and without microheater assisted degradation. Skin samples immediately after microneedle application (with and without microheater) were divided into three groups: Group 1 for tissue fluorescence imaging; Group 2 for histology processing and hematoxylin/eosin staining; and Group 3 for *in vitro* diffusion assay in PBS buffer.

For fluorescence imaging of tissue slide: rat skins were snap frozen in liquid nitrogen and embedded in O.C.T. Compound (Tissue-Tek) immediately to avoid fluorescence signal loss. Cryostat was performed to obtain skin sections in 5-µm thickness and stored at −20°C until imaging. Fluorescence microscopy of rat skin tissue slide (without mounting) was performed using a Nikon DS-Ri2 digital microscope camera mounted on a Nikon Eclipse E-600 microscope with a TRITC channel. Images were captured with the exact same exposure time and analog gain at 200× magnification using the NIS Elements Software (Nikon Corp.) following our published protocols.^[21]^

For histological staining, rat skin specimen was fixed in 4% paraformaldehyde and embedded in paraffin block for sectioning at 5-µm thickness. Skin slides were then subjected to hematoxylin/eosin staining as previously.^[22]^ Bright-field images at 200× magnification was taken to confirm successful insertion of microneedle with penetration of SC layers.

For *in vitro* diffusion assay, rat skin specimen after microneedle application (with and without microheater) were transferred into separate wells of 12-well plate containing 2 mL PBS buffer in each well to allow free floating of skin specimen. Small stir bars and stirring plate were used to ensure homogeneous distribution of Cy5 dye in the PBS solution. At various time points (10 min, 30 min, 1 h, 3 h, 6 h, 24 h, and 48 h) post incubation at room temperature, three aliquots of 50 µL PBS containing release Cy5 fluorescence dye were taken out while 150 µL fresh PBS were added back into the same well. Aliquots of dye-containing PBS were collected in black clear bottom 96-well plate and stored at 4°C until measurement. Following our previously published protocols,^[23]^ NIRF imaging of the 96-well plate containing time-dependent aliquots was performed with a filter pair of *Ex/Em* = 645/700 nm Identical illumination settings (auto exposure, medium binning, F/Stop = 2) were used for acquiring all NIRF images. The fluorescence intensity of each well was quantified with built-in function in Live Image software and presented as radiant efficiency [p/s/cm^2^/sr]/[µW/cm^2^] for each time point. Each experiment involving rat skin tests were repeated for at least three times.

## Acknowledgements

We are grateful for the financial support from NIH NIAMS R01AR064792, R21AR057512, R21AR072334, North American Spine Society (NASS), Virginia Microelectronics Consortium(VMEC), and University of Virginia Engineering in Medicine fund, Center for Advanced Biomanufacturing seed fund, start-up fund from Department of Orthopaedic Surgery at University of Virginia, and Commonwealth Health Research Board (CHRB) 207-10-18. We appreciate the histological support from Histology Core at University of Virginia.

## Supporting Information

**Supplementary Figure S1:**
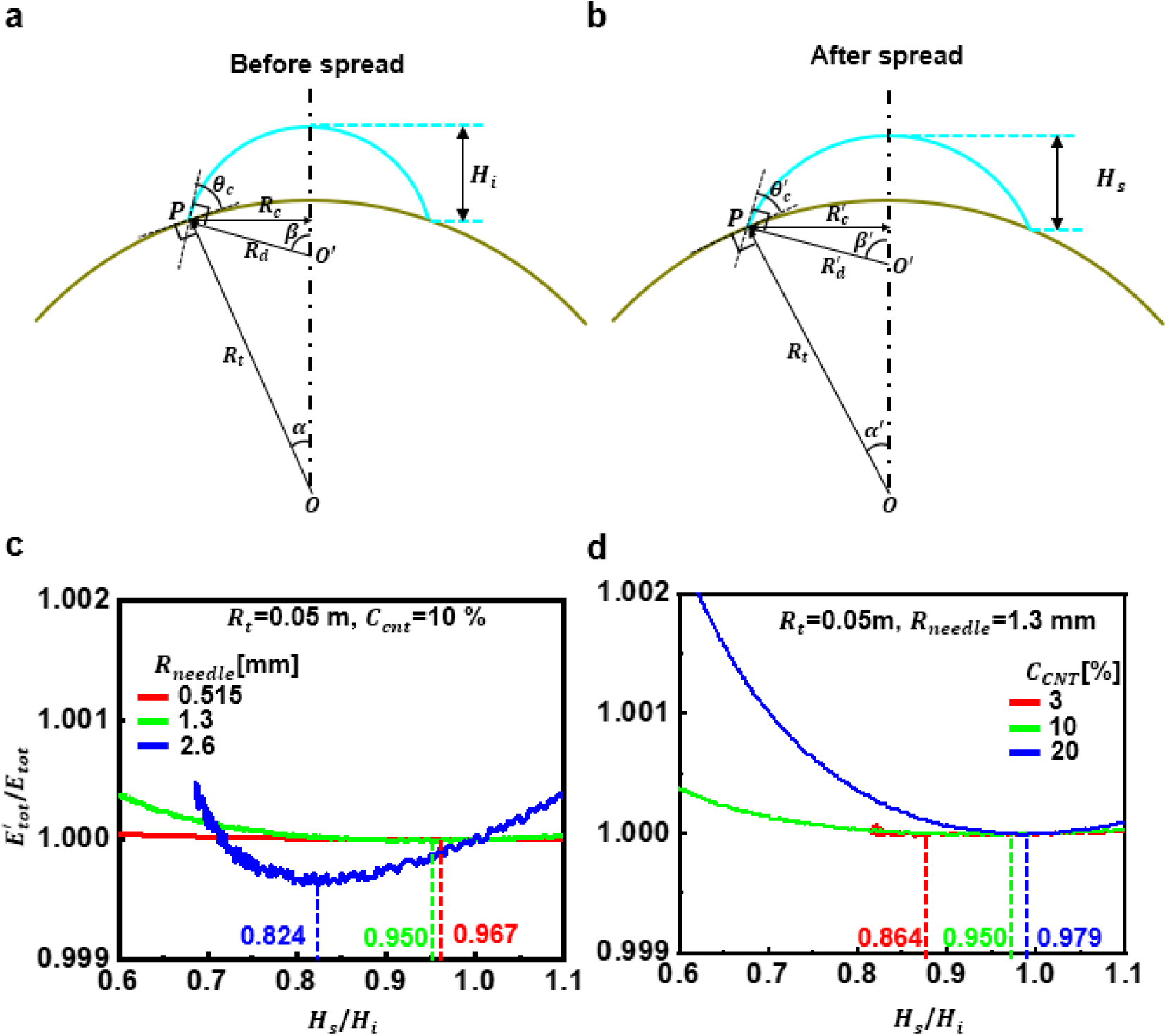
Schematic illustrations of the liquid droplet riding on a curved substrate before **(a)** and after **(b)** spreading. Effect of **(c)** the nozzle size of 3D printer and **(d)** the concentration of MWCNTs in MWCNTs/PDMS ink solution on the total energy after spreading normalized by the total energy before spreading 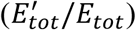. The minimization of 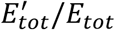 corresponds to the height of the droplet after spreading, *H*_*s*_*/H*_*i*_.

**Supplementary Figure S2:**
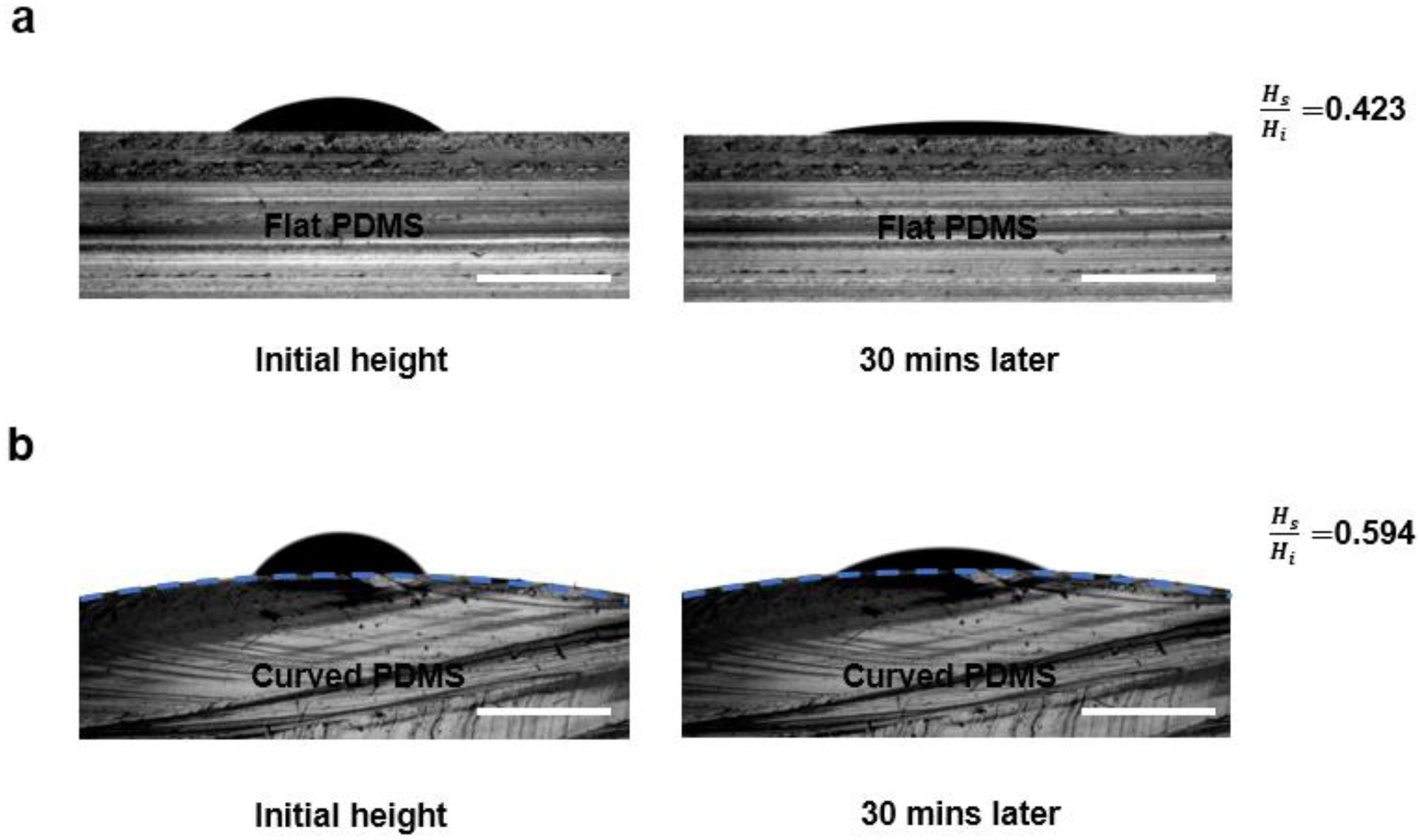
**(a)** Optical images of the MWCNTs/PDMS droplet (3% concentration of MWCNTs) on the flat PDMS substrate for as-printed and after 30-minute status. *Hs*/*Hi* = 0.423 indicates no spreading of droplet. **(d)** Optical images of the MWCNTs/PDMS droplet (3% concentration of MWCNTs) on the curved PDMS substrate (curve radius: 20 mm) for as-printed and after 30-minute status. *Hs*/*Hi* = 0.92 indicates no spreading of droplet. *Hs*/*Hi* = 0.594 indicates no spreading of droplet. All droplets in (c) and (d) were printed with the diameter of 18-gauge print head and were in 1.8 mm in diameter as printed. Scale bar is 2 mm.

**Supplementary Figure S3:**
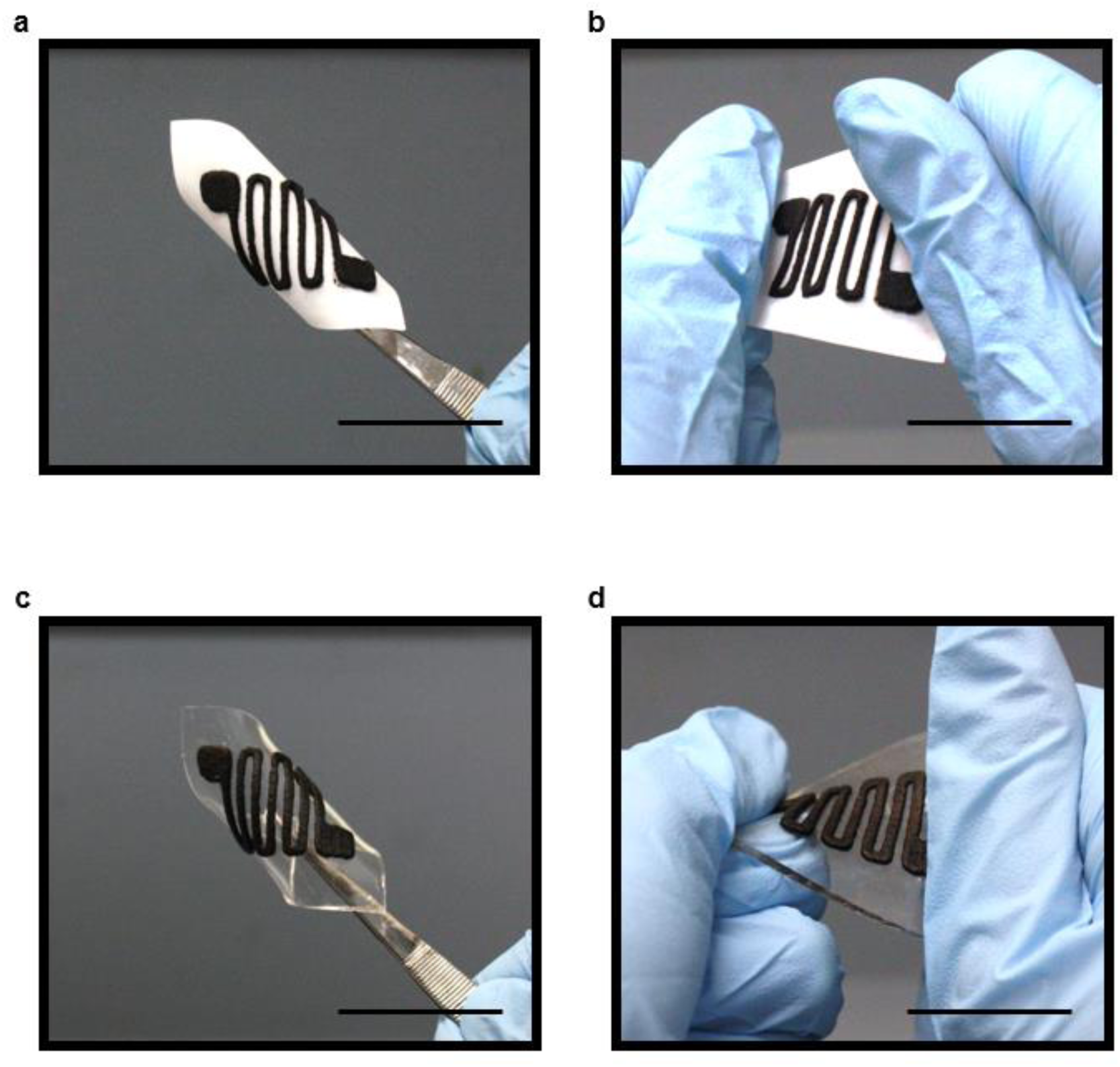
Optical images of. **(a)** Bended microheater on paper substrate. **(b)** Twisted microheater on paper substrate. **(c)** Bended microheater on PDMS substrate. **(d)** Twisted microheater on PDMS substrate. Scale bar is 30 mm.

**Supplementary Figure S4:**
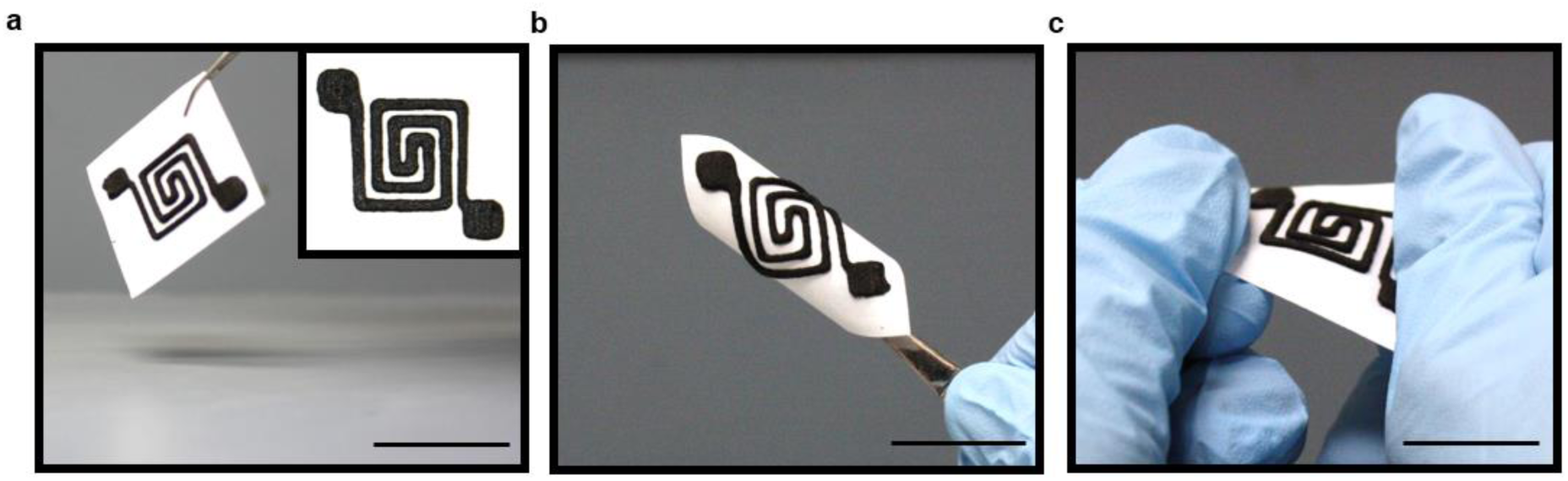
Optic images of: **(a)** Double spiral microheaters printed on paper substrate. **(b)** Bended microheater. **(c)** Twisted microheater. Scar bar is 35 mm for (a), and 25 mm for (b) and (c).

**Supplementary Figure S5:**
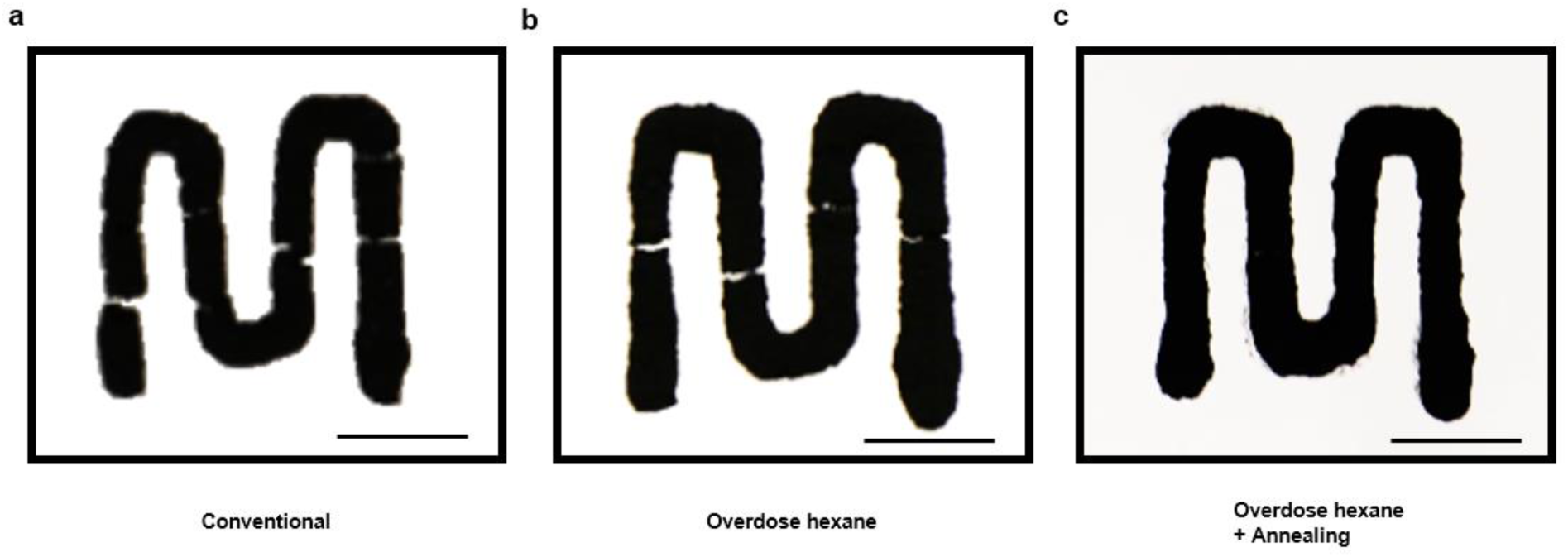
Optical images of printed simple pattern. **(a)** Pattern printed with conventional fabrication technique, i.e. MWCNTs/PDMS ink solution. **(b)** Pattern printed with overdose hexane added to MWCNTs/PDMS ink solution. **(c)** Pattern printed with overdose hexane and post-treated with annealing process. Scale bar is 3 mm.

**Supplementary Figure S6:**
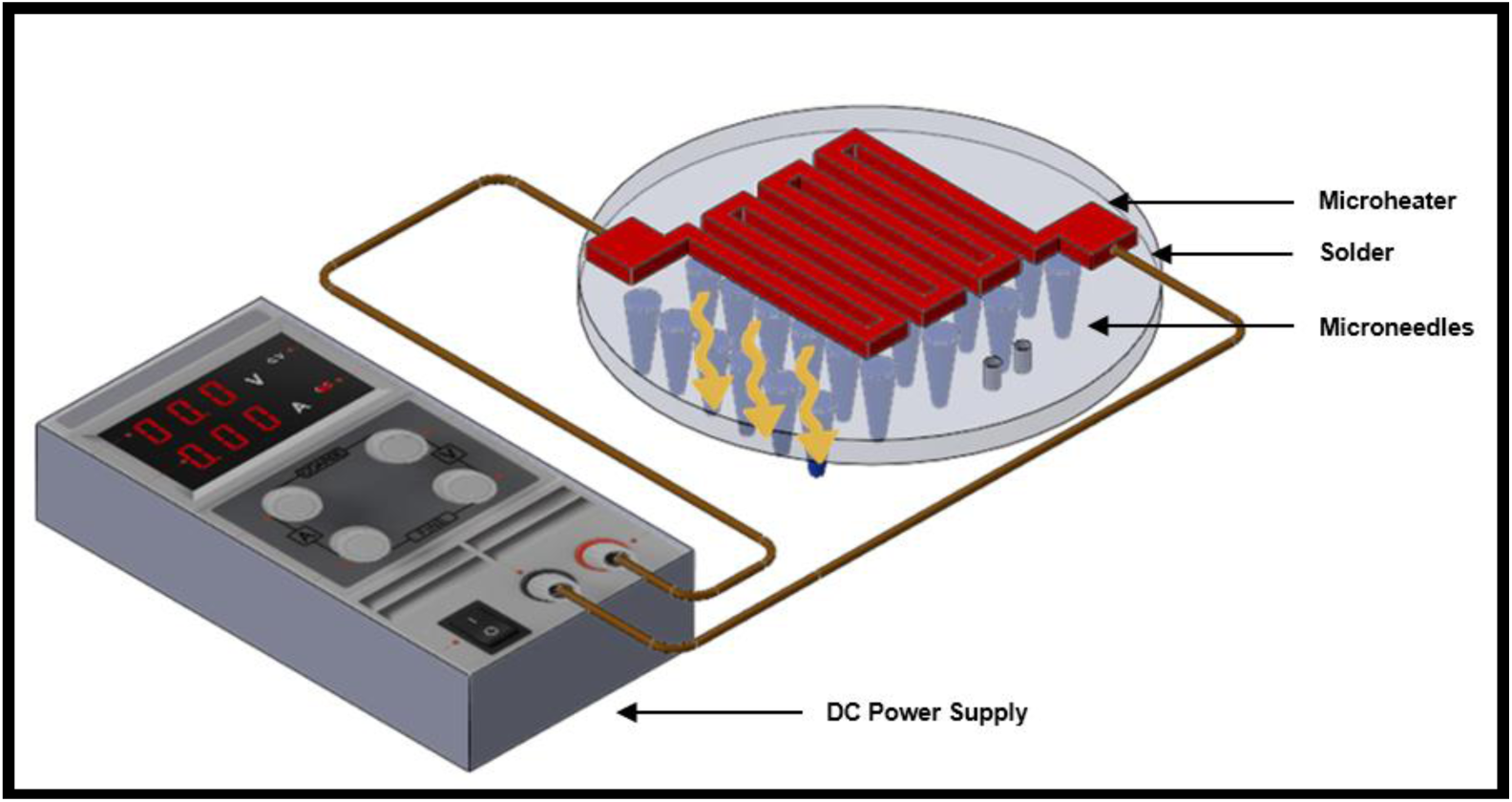
Schematic diagram of the Joule heating principle of microheater. A power supply is connected to both ends of microheater, and with the current flow in conductive composite, heat is generated due to Joule heating and transferred to patch, microneedle, and skins.

**Supplementary Figure S7:**
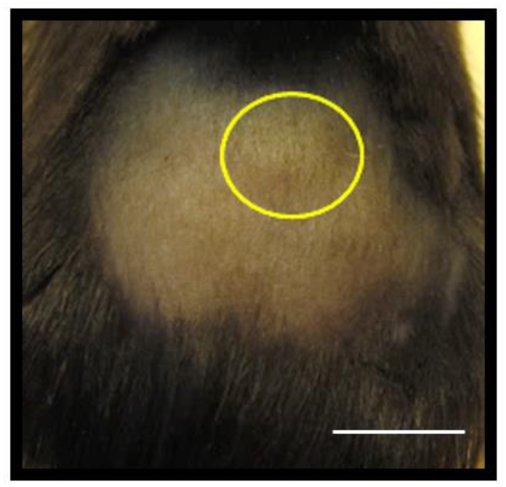
Optic Image of Live Mouse Dorsal Skin. Microheater/microneedle application did not cause visually detectable skin irritation on live mouse back skin. Yellow circle highlighted application site. No redness and skin rash etc. were observed after 24 hours of application. Scale bar is 20 mm.

**Supplementary Figure S8:**
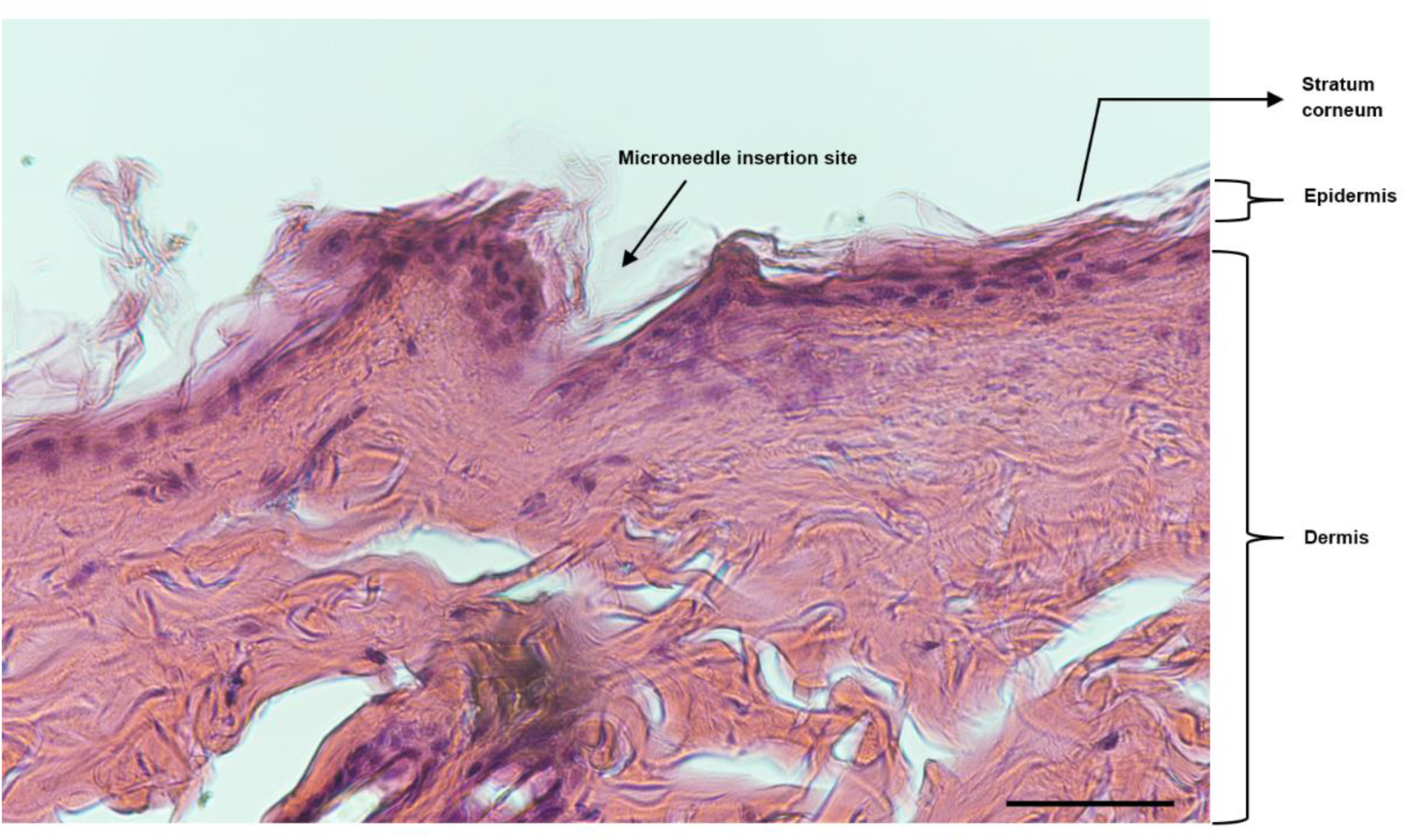
Representative histological image of hematoxylin-eosin staining of rat skin specimen confirmed the microneedle insertion and disruption of SC layer (×200 magnification). Scale bar is 100 μm.

